# Integrin α_6_ and EGFR Signaling Converge at Mechanosensitive Calpain 2

**DOI:** 10.1101/164525

**Authors:** Alyssa D Schwartz, Christopher L Hall, Lauren E Barney, Courtney C Babbitt, Shelly R Peyton

## Introduction

Carcinoma progression is associated with deposition of ECM that stiffens the local microenvironment [1, 2]. This tissue stiffening results in deposition of additional matrix proteins, initiating a positive feedback loop between cells and the evolving stroma [3]. Cells sense and respond to the stiffness of their environment via RhoA GTPase activation, which feeds back to increase cell contractility via activation of myosin light chain kinase [4]. These ECM-driven changes in cytoskeletal tension regulate motility in a cell-type specific manner [5]. *In vitro*, synthetic biomaterials have revealed that the type of material, stiffness, and biochemical surface modification alter the attachment and motility of cells [6, 7]. We therefore hypothesized that independently tuning stiffness, while altering integrin-binding sites in the presence of chemokinetic factors could reveal new mechanosensitive proteins that drive adhesion and motility in cancer cells.

Several mechanically responsive genes and proteins have been implicated in breast cancer metastasis [8]. For example, extracellular mechanical forces in the tumor microenvironment alter nuclear stiffness and gene expression [9] and activate adhesion proteins, including focal adhesion kinase (FAK) and talin [10-12], leading to increased motility. One class of surface receptors, integrins, translate extracellular forces to downstream signaling cascades in a process called *mechanotransduction*. Increasing substrate stiffness increases integrin binding and clustering, which has implications for several pathways in breast cancer metastasis, including FAK and PI3K signaling [13, 14]. However, most cancer mechanobiology research has focused on collagen- and fibronectin-binding integrins. Integrins that bind to other ECM proteins, including laminin investigated here, have largely been neglected, despite the prevalence of laminins, such as laminin-111 and −511, in the ECM of many tumor types, including breast and prostate cancers [15-17]. In the breast cancer cell line, MDA-MB-231, laminin-511 enhances cell adhesion and migration [18], while laminin-322 promotes cell survival [19].

EGFR has recently emerged as mechanosensitive [20, 21], and this process is critically relevant to cancer progression, given the known abundance of EGF in breast tumors [22] and frequent acquired resistance to EGFR inhibitors [23]. Residues on EGFR can be phosphorylated by α_V_β_3_ integrin clustering [24], so we postulated that there could be a role for integrins in facilitating EGFR mechanosensing [25].

To uncover whether such a relationship exists, we created a biomaterial system to identify mechanoresponsive proteins working cooperatively downstream of EGFR phosphorylation and laminin-binding integrin engagement in breast cancer. Through a combination of transcriptomics, molecular biology, and quantification of cell adhesion and motility, we found that the intracellular protease calpain 2 is one of these links. Gene expression quantification revealed that both calpain 2 and integrin α_6_ had an inverse relationship with ECM stiffness. Cell adhesion and motility experiments demonstrated that engagement of integrin α_6_ resulted in mechanosensitive effects on adhesion and motility in the same manner as EGF stimulation. This suggested coordination of integrin α_6_ with EGFR and that calpain 2 is downstream of both EGFR phosphorylation and integrin α_6_ engagement. Further, this indicated that calpain 2 is a common signaling node in the cell that regulates motility and cell adhesion in a mechanosensitive manner. In sum, we highlight the utility of tunable biomaterials systems to uncover previously unrealized relationships between different classes of proteins in breast cancer mechanobiology.

## Materials and Methods

### Cell Culture

All cell culture reagents were purchased from Thermo Fisher Scientific (Waltham, MA) unless otherwise specified. MDA-MB-231 cells were a generous gift from Sallie Schneider at the Pioneer Valley Life Sciences Institute. Highly metastatic and tropic MDA-MB-231 variants were kindly provided by Joan Massagué at the Memorial Sloan Kettering Cancer Center. These cell lines preferentially metastasize to the bone (1833 BoM [26]) brain (831 BrM2a [27]) or lung (4175 LM2 [28]). All cells were cultured in Dulbecco’s modified eagle medium (DMEM) with 10% fetal bovine serum (FBS) and 1% penicillin-streptomycin (P/S) at 37°C and 5% CO_2_.

### Synthesis of Poly(ethylene glycol)-phosphorylcholine (PEG-PC) Gels

PEG-PC gels were formed as previously described [29]. For this application, a 17wt% solution of 2-methacryloyloxyethyl phosphorylcholine (PC) (730114-5g, Sigma Aldrich, St. Louis, MO) in pH 7.4 phosphate buffered saline (PBS) was mixed with varying concentrations of poly(ethylene glycol) dimethacrylate (PEGDMA) (Sigma Aldrich). This solution was filtered through a 0.22 µm PVDF syringe filter, and excess oxygen was removed by degassing with nitrogen for 30 seconds. 20wt% Irgacure 2959 (BASF, Florham Park, NJ) was dissolved in 70% ethanol and added to the polymer solution in a ratio of 40 µL Irgacure to 1 mL of polymer solution. 50 µL of this final hydrogel precursor solution was sandwiched between a 3-(trimethoxysilyl)propyl-methacrylate treated 15mm diameter coverslip and an untreated coverslip and UV polymerized for 20 minutes. Gels were swelled in PBS for at least 48 hours before functionalizing the surface using 0.3 mg/mL Sulfo-SANPAH (c1111-100mg, ProteoChem, Hurricane, UT) in 50 mM HEPES buffer at pH 8.5, under UV for 10 minutes. Gels were then flipped onto droplets to achieve a theoretical final concentration of either 10 µg/cm² collagen 1 (A1048301, Thermo Fisher Scientific), 10 µg/cm² collagen 1 + 0.5 µg/cm² β_1_-chain-containing laminin isoforms (AG56P, Millipore,

Billerica, MA), or 10 µg/cm² collagen 1 + 20 ng/cm² EGF (R&D Systems, Minneapolis, MN), binding via accessible amines. Gels were incubated with protein overnight in a hydrated chamber at room temperature, washed in PBS, and sterilized under germicidal UV for 1 hour before cell seeding. For mechanical testing, gels were formed in a cylindrical mold (5µm high, 5µm diameter), allowed to swell for at least 48 hours and compression rheology was performed using an AR2000 (TA Instruments, New Castle, DE), as previously described [29].

### RNA-Seq

Total RNA was isolated using Gen Elute mammalian total RNA Miniprep kit (RTN70, Sigma Aldrich). The TruSeq stranded RNA LT kit (15032612, Illumina, San Diego, CA) was used to purify and fragment the mRNA, convert it to cDNA, and barcode and amplify the strands. Quality and length of the inserts was confirmed with an Agilent Genomics 2100 bioanalyzer, followed by single-end 75 bp reads on the MiSeq (Illumina) to generate a complete transcriptome from each sample. Transcripts were aligned to the hg19 human reference genome using the Tuxedo Suite pathway [30-33]. Cufflinks was used to determine statistically significant differential expression of genes (p<0.05) [34-36].

### qRT-PCR

Total RNA was isolated from two biological replicates as previously described [37]. 50 ng cDNA was then amplified using 10 pmol specific primers (Table S1) and the Maxima Sybr green master mix (Thermo Fisher Scientific) on a Rotor-Gene Q thermocycler (Qiagen, Valencia, CA) as follows: 50°C for 2 minutes, 95°C for 10 minutes followed by 45 cycles at 95°C for 10 seconds, 58°C for 30 seconds, and 72°C for 30 seconds. Both β-actin and ribosomal protein S13 were included as reference genes to permit gene expression analysis using the 2^-ddCt^ method.

### Cell Adhesion Quantification

MDA-MB-231 cells were seeded at a final density of 5,700 cells/cm^2^. Cells were either seeded immediately, or pretreated for 30 minutes before seeding with 20 µM ERK inhibitor FR180204 (Sigma Aldrich), 10 µg/mL calpain inhibitor IV (208724, Millipore), 20 µM lapatinib (LC Laboratories, Woburn, MA), or 100 ng/mL EGF. The plate was pre-incubated on the Zeiss Axio Observer Z1 microscope (Carl Zeiss AG, Oberkochen, Germany) for 1 hour, then cells were seeded onto gels functionalized with 10 µg/cm² collagen 1, 10 µg/cm² collagen 1 + 0.5 µg/cm² laminin, or 10 µg/cm² collagen 1 + 20ng/cm ^2^ bound EGF and imaging was started within 10 minutes of seeding. Images were taken with a 20x objective every 5 minutes for at least 2 hours. Cells were manually traced using Image J (NIH, Bethesda, MD). n ≥ 50 cells per condition.

### Cell Migration

Cells were seeded onto gels functionalized with 10 µg/cm² collagen 1 or 10 µg/cm² collagen 1 + 20ng/cm² EGF at a density of 5,700 cells/cm^2^ and allowed to adhere for 24 hours. For inhibitor conditions, cells were treated 2 hours prior to the start of imaging with 10 µg/mL calpain inhibitor IV, 20 µM ERK inhibitor FR180204, 20 µM lapatinib, or 10 ng/mL EGF (R&D Systems). Cells were imaged at 15 minute intervals for 12 hours on the Zeiss Axio Observer Z1 microscope (Carl Zeiss AG). n≥ 48 cells per condition were quantified using the manual tracking plugin for Image J (NIH, Bethesda, MD).

### Generation of ITGA6 and CAPN2 shRNA transduced cells

The shRNA sets specific to human ITGA6 (Sequence: CCGGCGGATCGAGTTTGAT AACGATCTCGAGATCGTTATCAAACTCGATCCGTTTTTG) and CAPN2 (sequence: CCGGCAGGAACTACCCGAACACATTCTCGAGAATGTGTTCGGGTAGTTCCTGTTTTTG) were purchased from Sigma-Aldrich (St. Louis, MO). Plasmids (in pLKO.1-puro) were packaged into virus particles according to manufacturer’s instructions and used to transduce MDA-MB-231-luciferase cells. Stable pools were selected using 1 mg/mL puromycin (Invivogen, San Diego, CA). Cells transduced with a non-targeting shRNA served as a control.

### Immunofluorescent Staining and Imaging

28,500 cells/cm^2^ were seeded and fixed at 24 hours in 4% formaldehyde. Cells were permeabilized in Tris-Buffered Saline (TBS) with 0.5% Triton-X and washed 3 times in TBS with 0.1% Triton-X (TBS-T). Blocking was done for 1 hour at room temperature in TBS-T + 2%w/v Bovine serum albumin (BSA, Sigma Aldrich) (AbDil). Cells were then incubated in primary antibody in AbDil for 1 hour at room temperature using 1 or more of the following antibodies; vinculin (V9264-200UL, 1:200, Sigma Aldrich), pEGFRY1068 (ab32430, 1:200, Abcam, Cambridge, MA), total EGFR (D38B1, 1:200, Cell Signaling, Danvers, MA), integrin α_6_ (ab134565, 1:200, Abcam), integrin α_3_ (ab131055, 1:200, Abcam), ERK (ab54230, 1:200, Abcam), calpain 2 (MABT505, 1:200, Millipore), or phalloidin 647 (A22287, Thermo Fisher Scientific). For insoluble fractions, cells were permeabilized in cold Tris-Buffered Saline (TBS) with 0.5% Triton-X at 4°C for 1 minute prior to fixing. For immunofluorescent imaging of ECM proteins, cells were fixed at 24 hours (57,000 cells/cm^2^) or 6 days (11,400 cells/cm^2^) before staining with a pan-laminin antibody (ab11575, 1:100, Abcam), collagen 1 antibody (ab6308, 1:200, Abcam), or Fibronectin-488 antibody (563100, 1:200, BD Biosciences, San Jose, CA). Molecular Probes secondary antibodies were used at a 1:500 dilution (goat anti-mouse 555 (A21422) and goat anti-rabbit 647 (A21244) or goat anti-rabbit 488 (A11008), Thermo Fisher Scientific). Cells were then treated with DAPI at a 1:5,000 dilution for 5 minutes, and washed in PBS prior to imaging of 2 biological replicates on a Zeiss Cell Observer Spinning Disk (Carl Zeiss AG).

### Matrix Decellularization

Cells were cultured on PEG-PC gels of 1, 4, 8, and 41 kPa for 24 hours or 6 days, and the resulting matrix created by the cells was visualized by adapting a previously published protocol [38]. Briefly, cells were washed with warm PBS, then lysed with 500µL warm extraction buffer (PBS with 0.5% Triton-X with 20 mM ammonium hydroxide) for 10 minutes at room temperature. 1 mL of PBS was added and the gels were stored overnight at 4°C before staining as described above.

### Protein and Phospho-protein Quantification

Autolyzed calpain 2 was analyzed via western blot. Cells were cultured on PEG-PC gels of 1 or 41 kPa for 24 hours, then lysed in RIPA buffer with protease inhibitors: 1 cOmplete Mini EDTA-free Protease inhibitor cocktail tablet per 10 mL (Roche, Indianapolis, IN) 1 mM phenylmethylsulfonyl fluoride (Thermo Fisher Scientific), 5 μg/mL pepstatin A (Thermo Fisher Scientific), 10 μg/mL of leupeptin (Thermo Fisher Scientific), and phosphatase inhibitors: Phosphatase inhibitors cocktail-II (Boston Bioproducts, Boston, MA), 1 mM sodium pyrophosphate (Thermo Fisher Scientific), 25 mM β-glycerophosphate (Santa Cruz, Dallas, TX). Lysates were prepared in a 5x reducing sample buffer (39000, Thermo Fisher Scientific), run on Tris-Glycine gels in an Invitrogen mini-Xcell Surelock system, and then transferred to nitrocellulose membranes (88018, Life Technologies, CA). Membranes were blocked with AbDil for 1 hour, then stained overnight at 4°C with antibodies against calpain 2 (MABT505, 1:1000, Millipore), integrin α_6_ (ab134565, 1:2000, Abcam), EGFR (D38B1, 1:1000, Cell Signaling Technology, Danvers, MA), integrin α_3_ (343802, 1:1000, Biolegend, San Diego, CA), integrin β_4_ (ab29042, 1:1000, Abcam), GAPDH (ab9485, 1:2000, Abcam) or β-actin (ab75186, 1:1000, Abcam). Membranes were washed 3 times with TBS-T, stained in secondary antibody (Goat anti-rabbit 680 (926-68021, 1:20,000) and Donkey anti-mouse 800 (926-32212, 1:10,000), LiCor, Lincoln, NE) for 1 hour at room temperature, protected from light, and washed 3 times in TBS-T with a final wash in TBS. Membranes were imaged on the Odyssey CLx (Licor).

2 biological replicates of lysates were collected during cell adhesion at time 0, at 5 minutes, and at 24 hours. At 5 minutes, the adhered fraction was lysed off the surface, while suspended cells were pelleted, then lysis buffer was added. These 2 fractions were combined for analysis. Phospho-protein levels were quantified with the MAGPIX system (Luminex, Austin, Texas) with MILLIPLEX MAP Multi-Pathway Magnetic Bead 9-Plex - Cell Signaling Multiplex Assay (48-681MAG, Millipore) and added beads against p-EGFR (pan-tyrosine, 46-603MAG, Millipore), p-MEK (Ser222, 46-670MAG, Millipore), and p-ERK1/2 (Thr185/Tyr187, 46-602MAG, Millipore), following the manufacturer’s protocols, with beads and antibodies used at 0.25x.

### Co-Immunoprecipitation

MDA-MB-231 cells were grown to confluence in a T-75 cell culture flask. Cells were treated with 10 mL of fresh media containing 10% FBS, with or without an additional 100 ng/mL EGF for 10 minutes. Flasks were lysed in 4.5 mL of lysis buffer, comprised of 50 mM Tris (pH 7.4), 0.15 M NaCl, 0.1% Triton-X, with protease and phosphatase inhibitors as described above. 1 mL of lysate was immunoprecipitated with 10 µL agarose beads (Thermo Fisher Scientific) with either 5 µL rabbit IgG (ab6718, Abcam), 10 µL anti-EGFR (D38B1, Cell Signaling), or with 10 µL anti-EGFR pre-conjugated to sepharose beads (5735S, Cell Signaling). Lysates were precipitated on beads overnight at 4°C on a rotating platform and spun down. Beads (with protein) were washed 3x with TBS, and boiled in 70 µL of DiH_2_O and sample buffer. Blotting was done as described above for pEGFR (ab32430, Abcam) and integrin α_6_ (ab134565, Abcam).

### Statistical Analysis

One-way Analysis of Variance (ANOVA) with a Tukey post-test was performed on paired samples using Prism v5.04 (Graphpad software, La Jolla, CA). Data reported is the mean and reported error is standard deviation with significance values of p ≤ 0.05 as *, p ≤ 0.01 as **, and p ≤ 0.001 as ***.

## Results

### Cells Maximize Expression of Integrin α_6_ and Extracellular Laminin on Soft Substrates

To identify ECM and integrin-associated genes involved in breast cancer mechanosensing, we performed whole transcriptome sequencing (RNA-Seq) on MDA-MB-231 cells cultured on collagen 1-functionalized PEG-PC hydrogels ranging in stiffness from 1 to 41 kPa for either 24 hours or 6 days (Fig. S1a). The stiffness range chosen spans that of several tissues in the body where breast cancer metastasizes (1 kPa- brain [39], 4 kPa- bone marrow [40], 8 kPa- lung [41]), as well as a supra-physiological stiffness (41 kPa) that approximates extremely stiff breast tumors [42] and a tissue culture polystyrene (TCPS) control (Fig. S1).

Because the ECM is known to regulate the ability of cancer cells to migrate and metastasize, we focused our in-depth analysis of this RNA-Seq data set on genes that regulate interaction with the ECM. We found that the expression of integrins, other focal adhesion complex genes, and genes downstream of EGFR phosphorylation became sensitive to stiffness over time (Fig. 1a-b, Fig. S1b). Short-term (24 hour) decreases in expression were clear in integrin α_6,_ calpain 2, and integrin β_4_ (Fig. 1a). However, integrin β_4_ expression was 3-fold higher on gels than on TCPS after 6 days of culture (Fig. 1b), which we confirmed at the protein level (Fig. S2a). This implies a switch in dimer pairs, as integrin α_6_ preferentially dimerizes with integrin β_4_ over integrin β_1_ [43]. This is an important finding, as α_6_β_4_ is known to be important in cancer cell motility and invasion [44]. We compared the day 6 gene expression data from Figure 1b to that at 24 hours from Figure 1a and found that the expression of this subset of genes was generally higher at day 6 (Fig. S1b), suggesting a time-dependent mechanosensing event related to this set of integrins, focal adhesion proteins, and EGFR.

**Figure 1.**
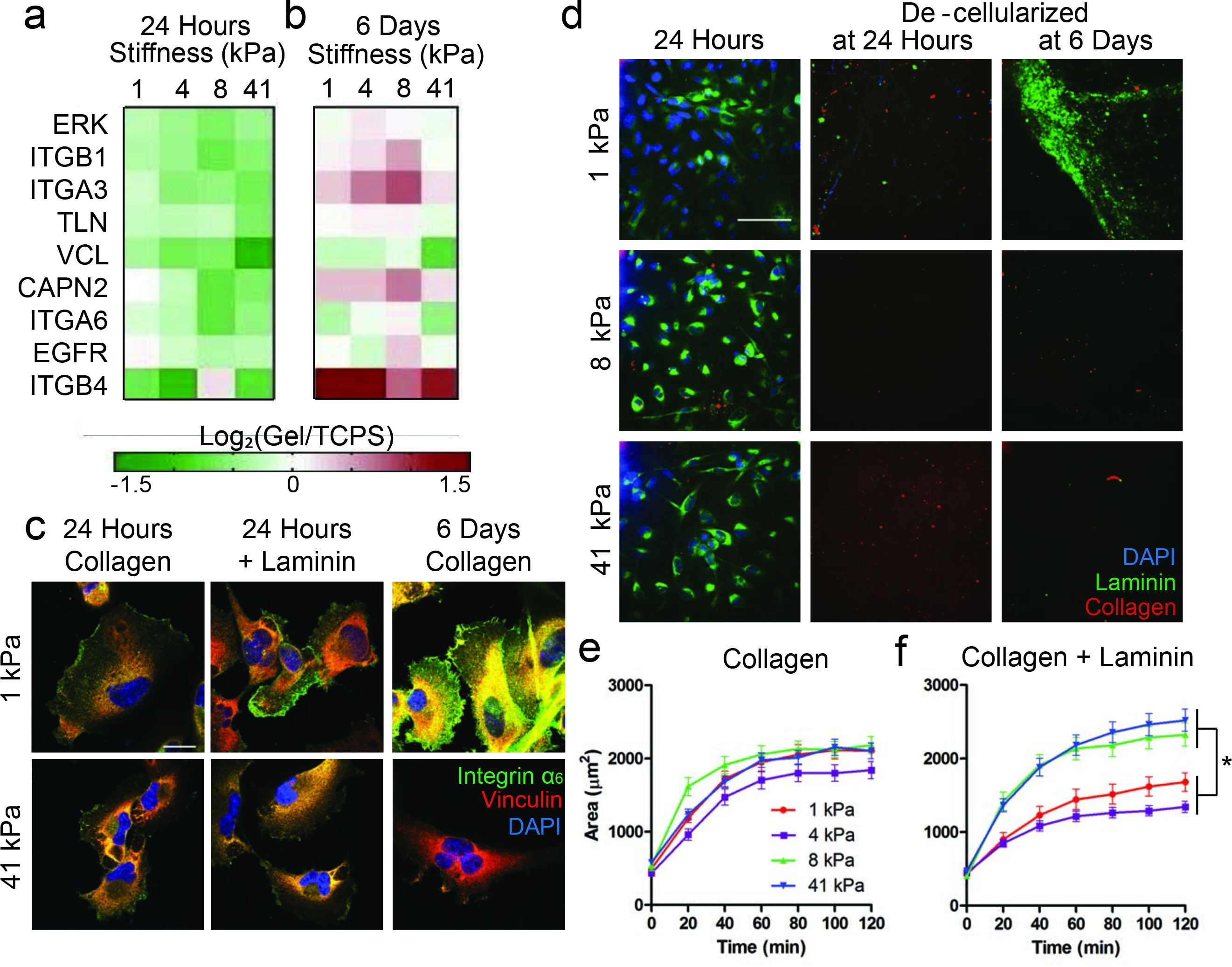
Cells on soft substrates have increased integrin α_6_ expression and produce extracellular laminin. (a-b) Log_2_(fold change) of gene expression from cells cultured on gels of stiffnesses 1, 4, 8 or 41 kPa relative to expression on TCPS of genes after 24 hours (a) or 6 days (b), determined through qRT-PCR. Decreased expression = green and increased expression = red. N=2. (c) Representative immunofluorescent staining of vinculin (red) and integrin α_6_ (green) and DAPI (blue) in cells cultured on gels with collagen only for 24 hours, collagen with laminin for 24 hours, or collagen only for 6 days. Yellow denotes overlap of vinculin and integrin α_6_ staining. Scale bar = 20 µm. (d) Representative Immunofluorescent staining for ECM proteins after 1 day of culture (left), decellularized gels after 24 hours (center) or decellularized gels after 6 days (right). Collagen 1 = red, pan-laminin = green, DAPI = blue. Scale bar = 100 µm. (e-f) Cell area during adhesion quantified on 1, 4, 8, and 41 kPa gels functionalized with collagen 1 only (e) or collagen 1 + laminin (f). N=3 biological replicates, n=75 cells per condition.

Given that integrin α_6_β_4_ is laminin-binding, and that we saw mechanosensitive changes in α_6_ gene and protein expression (Fig. S2b), we hypothesized that we could tune integrin α_6_ mechanosensing on these surfaces by supplementing the collagen-functionalized hydrogel surfaces with β_1_-chain-containing laminins. On soft gels, integrin α_6_ expression was higher at 24 hours with added laminin compared to collagen 1 only, and it was also high after 6 days of culture, even in the absence of supplemented laminin (Fig. 1c, Fig. S2c). This mechanosensitivity appears to be specific to integrin α_6_ as other laminin-binding integrins (e.g. integrin α_3_) did not follow this trend (Fig. S2d-e).

As we noted this integrin α_6_ effect when we supplied only collagen 1, we hypothesized that cells might be producing their own laminin to support integrin α_6_ binding at the longer culture times. Immunofluorescent staining revealed that laminin, but not collagen 1 (Fig. 1d) or fibronectin (Fig. S3a), was upregulated on the softer gels at 6 days. We separately confirmed that differential staining was not due to ECM proteins crosslinked to the gels or from serum (Fig. S3b). Cultures that were decellularized after 6 days showed the highest amount of extracellular laminin on the softest, 1 kPa gels (Fig. 1d), concordant with the increase in integrin α_6_ expression in Figure 1c.

To understand the impact of laminin and integrin α_6_ expression on cell phenotype, we quantified cell spreading during initial adhesion to the hydrogel surfaces. We found cells spread to similar extents across the 4 stiffnesses examined here when only collagen 1 was present (Fig. 1e). However, when laminin (0.5µg/cm^2^) was added to the collagen (10µg/cm^2^) on the gel surfaces, cell spreading was mechanosensitive (Fig. 1f). We attempted to remove collagen from the experiment completely, but found that cells did not adhere to laminin-only hydrogels (data not shown). This was not surprising given the documented role of laminin being somewhat anti-adhesive [45]. In sum, cells upregulate integrin α_6_ expression on soft substrates, which correlates with secretion of extracellular laminin. This coordination appears to take several days, but the mechanosensitivity effect on cell spreading can be accelerated by adding β_1_-chain-containing laminin to the culture substrata.

### Cell Response to EGF Stimulation Mirrors Mechanosensing on Laminin

Given the prevalence of EGF in the tumor microenvironment and its known role in promoting cancer cell motility, we posited that the mechanosensitive expression of EGFR alongside the laminin-binding integrin α_6_ implied that they mechanotransduce under a related pathway. Similar to previous reports, we found that MDA-MB-231 cells spread to a smaller area on hydrogels upon addition of EGF prior to and during adhesion [37], achieving a similar size to cells on soft, laminin-coated gels without EGF (Fig. 2a). EGFR is known to associate with the α_6_β_4_ integrin heterodimer to enhance clustering and response to EGF [46], and in agreement with this report, we found that the addition of EGF increased the association of integrin α_6_ with EGFR in our system (Fig. S4a). We separately analyzed the protein expression of integrin α_6_ and EGFR in variants of the MDA-MB-231 cell line selected for their ability to metastasize specifically to the brain, bone, and lung [26-28]. The brain tropic cells, which metastasize to the softest of these sites, have the highest expression of both integrin α_6_ and EGFR (Fig. S4b).

**Figure 2.**
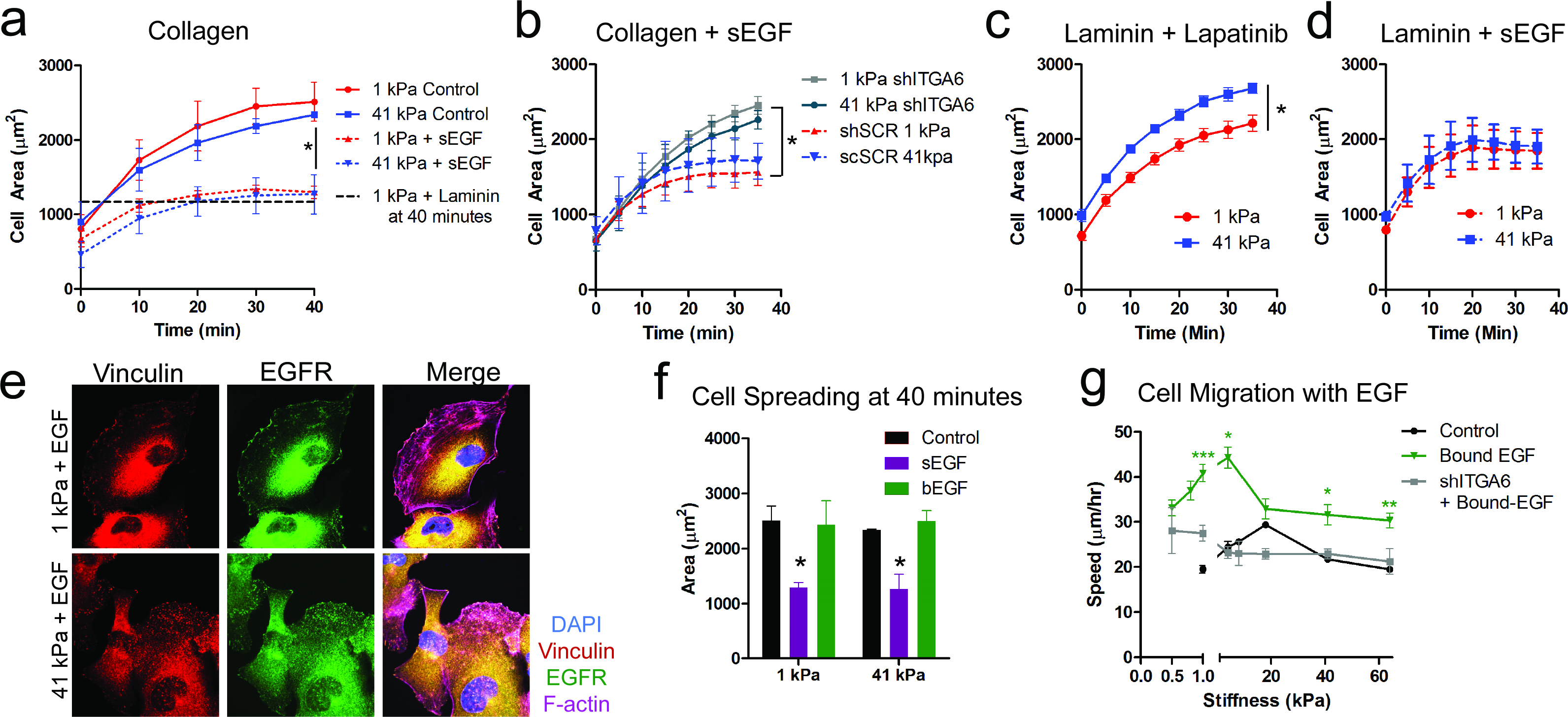
Addition of EGF mimics mechanosensing on soft, laminin containing surfaces. (a) Cell area during adhesion with soluble EGF compared to data from Fig. 1d-e. N=2, n=50. (b) Cell area during adhesion of integrin α_6_ knockdown cells (shITGA6) compared to scramble control cells (shSCR). N=2, n=50 cells. (c) Cells were pretreated with lapatinib for 30 minutes prior to seeding onto 1 and 41 kPa gels functionalized with collagen 1 and laminin and traced to quantify cell area over time. N=2, n=50 cells. (d) Cells were pretreated with EGF for 30 minutes prior to seeding onto 1 and 41 kPa gels functionalized with collagen 1 and laminin and traced to quantify cell area over time. N=2, n=50 cells. (e) Representative images of cells seeded onto gels of 1and 41 kPa functionalized with collagen 1 and EGF stained for vinculin (red), EGFR (green), F-actin (purple), and DAPI (blue). (f) Area was quantified 40 minutes after adhesion to 10 µg/cm^2^ collagen 1 with no EGF, soluble EGF (sEGF), or bound EGF (bEGF). Statistics are shown relative to the control bar. N=2, n=50 cells. (g) Parental MDA-MB-231 cell speeds were quantified on 10 µg/cm^2^ collagen 1 with no EGF, with 10 ng/mL soluble EGF, 20 ng/cm² bound EGF. shITGA6 cell speeds were quantified on 10 µg/cm^2^ collagen 1 + 20 ng/cm² bound EGF. N=2, n=48 cells. Asterisks denote significance (p<0.05) relative to the no EGF control at the same stiffness.

To test the necessity of integrin α_6_ in the observed response to EGF, we generated a stable shRNA knockdown cell line (shITGA6, Fig. S4c-e). We quantified adhesion of the shITGA6 cells to collagen-functionalized gels, and found that they spread to a larger area during adhesion with EGF stimulation, compared to cells transfected with the scramble control (shSCR), suggesting a role for integrin α_6_ during EGF response on collagen (Fig. 2b). Even though there was no laminin present, others have demonstrated a role for integrin α_6_ in facilitating growth factor response, independent of laminin binding [47]. Combined with our data, this implicates a mechanosensitive role for integrin α_6_ during EGFR signaling (Fig 2b).

### EGF Overwhelms Laminin-Based Mechanosensing and Maximizes Cell Motility on Soft Substrates

To further explore the relationship between integrin α_6_ and EGFR signaling during adhesion, we functionalized hydrogels with collagen 1 and laminin, and treated the cells with lapatinib, a dual tyrosine kinase inhibitor against EGFR and HER2. This approach allowed cells to engage integrin α_6_ during adhesion, while preventing EGFR signaling by obstructing ATP-binding on the intracellular domain of EGFR and HER2. When pretreated with lapatinib, we found that cells adhering on 41 kPa hydrogels spread to a significantly larger area than cells on 1 kPa hydrogels (Fig. 2c). This result mimics the adhesion we observed of cells on laminin without supplemented EGF or lapatinib (Fig. 1f). Further, when we supplemented cells with soluble EGF on the surfaces functionalized with collagen 1 and laminin, cells on both 1 and 41 kPa spread to a smaller area (Fig. 2d). This EGF-induced reduction in spreading is similar to our observations of cells on collagen 1 only in the presence of soluble EGF (Fig. 2a). Together, these data suggest that, even in the presence of laminin, adhesion signaling through EGFR dominates, but in the absence of exogenous EGF, signaling via mechanosensitive integrin α_6_ controls cell spreading during adhesion.

We observed that sEGF caused rapid internalization of EGFR, within 10 minutes of administration (Fig. S4f). To promote sustained EGFR signaling to examine its relationship with integrin α_6_ [46], we covalently linked both collagen 1 and EGF to the surface of the hydrogels. This EGF presentation minimized EGFR internalization (Fig. S4f) and localized both pEGFR(Y1068) and total EGFR to regions of vinculin staining (Fig. 2e, Fig. S5a). EGFR receptor internalization was required for EGF to moderate adhesion, as only cells supplemented with soluble EGF showed measurable differences in cell area after 40 minutes (Fig. 2f). However, bound EGF had a measurable effect on cell motility (Fig. 2g). First, we observed a biphasic trend in cell speed with respect to stiffness. This phenomenon has previously been described in many cell types, including smooth muscle cells [7] and glioma cells in both 2D [48] and 3D [49]. In this migration study, only collagen 1 was present, suggesting that collagen-binding integrins contributed to this mechanosensitive motility after 24 hours [50]. Our data agree with literature reports demonstrating that cells seeded on collagen 1 exhibit mechanosensitive motility [51], but do not differ in adherent area between 1 and 30 kPa [52]. The addition of bound EGF increased cell motility across the stiffness range tested here (Fig. 2g), and shifted the biphasic curve to softer substrates, requiring us to expand the mechanical range of the hydrogels to capture the full curve (Fig. S5b) [29]. We also observed a role for integrin α_6_ in regulating this EGF-dependent motility, as shITGA6 cells were less responsive to EGF than the control (Fig. 2g). The scramble control cells, which peaked at 34 µm/hr on 1 kPa gels, were more responsive to EGF than the shITGA6 cells (Fig. S5c). Interestingly, shITGA6 cells don’t respond to changes in stiffness from 1 to 18 kPa, either in control media, or in response to soluble EGF (Fig. S5d). Together, this data suggests that the addition of EGF alters mechanosensitive motility and that the MDA-MB-231 breast cancer cells are dependent on integrin α_6_ for this response.

### Calpain 2 is Downstream of both EGFR and Integrin α_6_ Mechanosensing

Inspired by the trends observed in the RNA-seq data (Fig. S1), we clustered our PCR data from the 24 hour time point, which uncovered that ITGA6 and EGFR expression clustered with the intracellular protease CAPN2 (Fig. 3a). Calpain 2, which has been implicated in focal adhesion formation and disassembly, is thought to play an important role in breast cancer cell motility and metastasis [53]. Because calpain 2 is activated by ERK [54], we performed phospho-analysis of multiple kinases in integrin- and EGFR-dependent pathways (Fig. S6a). Steady-state phosphorylation of EGFR, MEK, and ERK increased on stiffer surfaces, as previously demonstrated [20, 55, 56] (Fig. S6b). However, we observed the opposite trend with the activation of calpain 2 (Fig. S6c).

**Figure 3.**
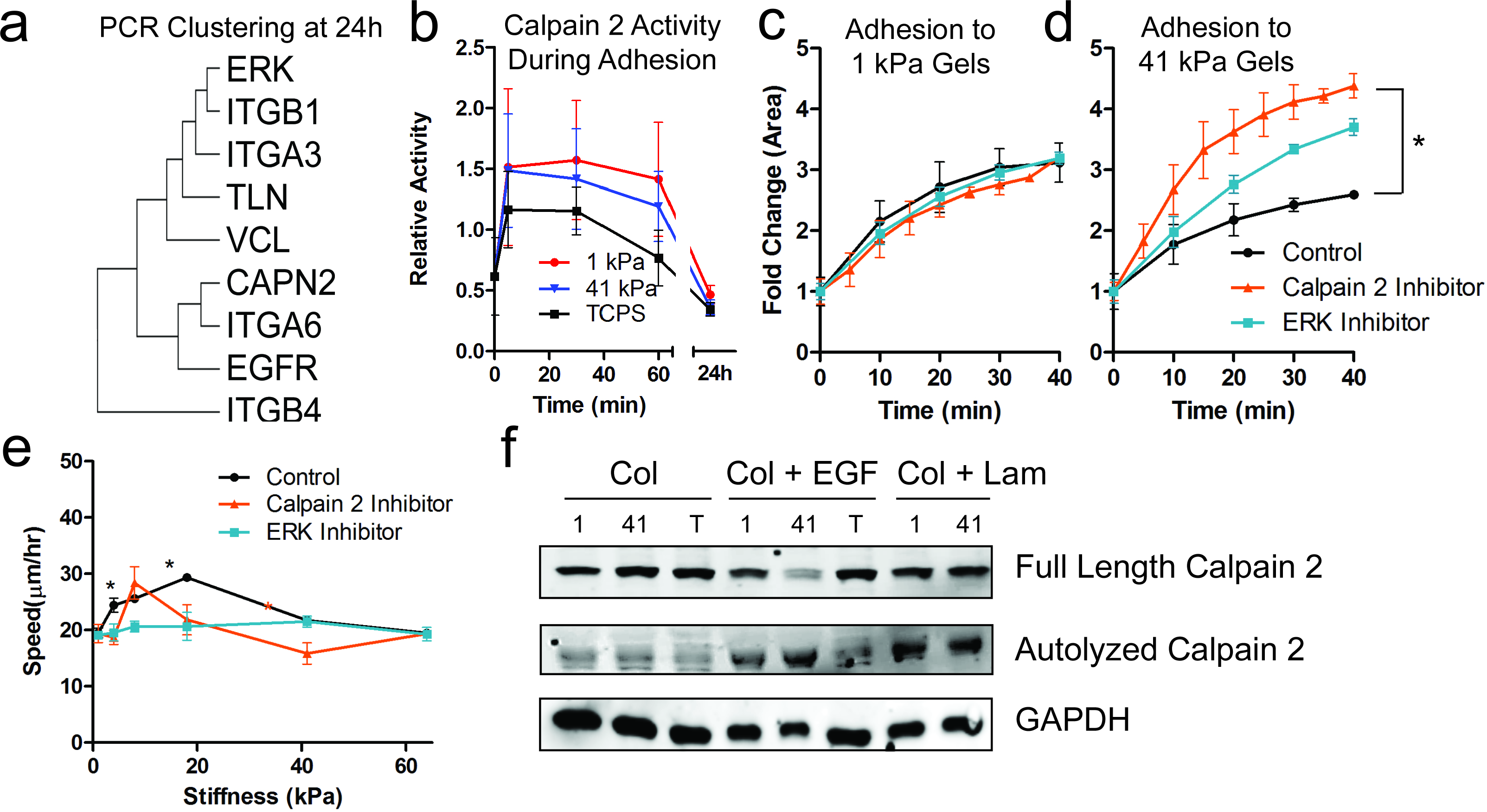
Calpain 2 is downstream of both mechanosensitive integrin α_6_ and EGFR. (a) Clustering of PCR data on genes expressed after 24 hours of culture on gels functionalized with 10µg/cm^2^ collagen 1 (data in Fig. 1a). (b) Calpain 2 activity during adhesion was measured by evaluating the ratio of autolyzed/full length calpain 2 via western blot in both the adhered and suspended cell fractions during the first 60 minutes, or just the adhered fraction after 24 hours. N=2. (c-d) Cell area was quantified during adhesion in the presence of an ERK1/2 inhibitor or calpain 2 inhibitor on 1 kPa (c) and 41 kPa (d) gels. N=2, n=50. (e) Cell migration speeds quantified the presence of an ERK1/2 or calpain 2 inhibitor. N=2, n=48. (f) Representative western blot for full length and autolyzed calpain 2, with loading control GAPDH. In order of lanes from left to right: 1 kPa with collagen 1, 41 kPa with collagen 1, TCPS, 1 kPa with collagen 1 + 100ng/mL soluble EGF for 10 minutes, 41 kPa with collagen 1 + 100ng/mL soluble EGF for 10 minutes, TCPS + 100ng/mL soluble EGF for 10 minutes, 1 kPa with collagen and laminin, 41 kPa with collagen and laminin.

We observed higher calpain 2 activity (autolysis) during adhesion to the soft hydrogels compared to tissue culture plastic (Fig. 3b). Although repeated multiple times, this trend was not statistically significant. There was a functional role for calpain 2 during mechanosensing on the gels, however, as a pharmacological inhibitor to either calpain 2 or ERK had no effect on cell spreading on 1 kPa gels, but resulted in significantly larger cells on 41 kPa gels (Fig. 3c-d). In parallel, to provide evidence of the necessity of calpain 2 in responding to EGF stimulation, we generated a CAPN2 knockdown cell line (Fig. S7a-c), and observed that those cells were less responsive to EGF than the scramble control cells during adhesion, independent of stiffness (Fig. S7d). Further, calpain 2 inhibition was sufficient to limit cell response to soluble EGF using a pharmacological inhibitor (Fig. S7e). However, when we limited calpain 2 activity in integrin α_6_ knockdown cells, the effect of larger cells during adhesion was exaggerated (Fig. S7e).

We further hypothesized that altering calpain 2 activity could disturb the typical biphasic relationship between cell migration speed and matrix stiffness given its known role in mediating focal adhesion turnover [54]. Inhibiting ERK activity entirely eliminated the durokinesis effect, and calpain 2 inhibition shifted the migration curve maximum to lower stiffnesses (Fig. 3e). The limiting step during motility on stiff surfaces is retraction of the rear edge [57], and here we demonstrate that inhibiting calpain 2 reduces cell migration on stiffer substrates, likely because of this reduced ability to turn over adhesions. We then compared migration speeds of the integrin α_6_ knockdown and scramble control cells from figure 2, in response to the calpain 2 inhibitor (Fig. S5d). We found that the scramble control cells responded to the inhibitor similarly to the parental cells, although the addition of soluble EGF was able to partially overcome that inhibition. However, the shITGA6 cells exhibited slower migration at 18 kPa even in the control media, so the calpain 2 inhibitor had no additional effect (Fig. S5d).

After 24 hours of culture on PEG-PC gels with collagen 1 or collagen 1 and bound EGF, cells adopted a normal morphology with no noticeable protrusions (Fig. S7f). However, in the presence of lapatinib, ERK inhibitor, or calpain 2 inhibitor, we observed that cells left behind long, stable protrusions at the rear of the cell (Fig. S7f), indicative of their inability to release adhesion sites [58]. There were minimal observable differences in morphology as a function of stiffness, regardless of the pharmacological treatment applied. This suggests that at these concentrations, the inhibitors impact focal adhesion turnover enough to functionally alter motility with marginal impacts on morphology at these stiffnesses.

While we demonstrated earlier that integrin α_6_ is necessary for both mechanosensitive and EGFR-mediated adhesion, this data suggests that EGFR activates calpain 2 to regulate mechanosensitive motility in breast cancer cells. To some extent on collagen-functionalized hydrogels, and particularly on plastic, calpain 2 activity was low, and EGF stimulation increased calpain 2 activity on the hydrogels (Fig. 3f). Finally, when we supplemented the gels with laminin, calpain 2 activity was maximized on both stiffness gels, suggesting an additional role for integrin α_6_ engagement to activate calpain 2. Interestingly, when integrin α_6_ is knocked down in a stable manner, the cells still exhibit calpain 2 activity, suggesting that the cells might be compensating for the lack of integrin α_6_ signaling via another mechanism, not explored here (Fig. S7g). Together, these data suggest that the mechanosensitive receptors integrin α_6_ and EGFR both activate calpain 2 to facilitate stiffness-dependent adhesion and migration.

## Discussion

This work demonstrates integrin α_6_ and calpain 2 as mechanosensitive proteins important for stiffness-driven breast cancer cell adhesion and migration (Fig. 4a). The mechanosensitivity of calpain 2 is a new finding, as well as the role of integrin α_6_ in EGFR signaling that facilitates cell adhesion (Fig. 4b). EGF is known to enhance breast cancer cell adhesion and motility, and here we demonstrate that mechanosensitive integrin α_6_ and EGFR both signal through calpain 2. We also contribute that in the absence of EGF stimulation, integrin α_6_ and calpain 2 regulate stiffness-dependent adhesion and motility (Fig. 4b).

**Figure 4.**
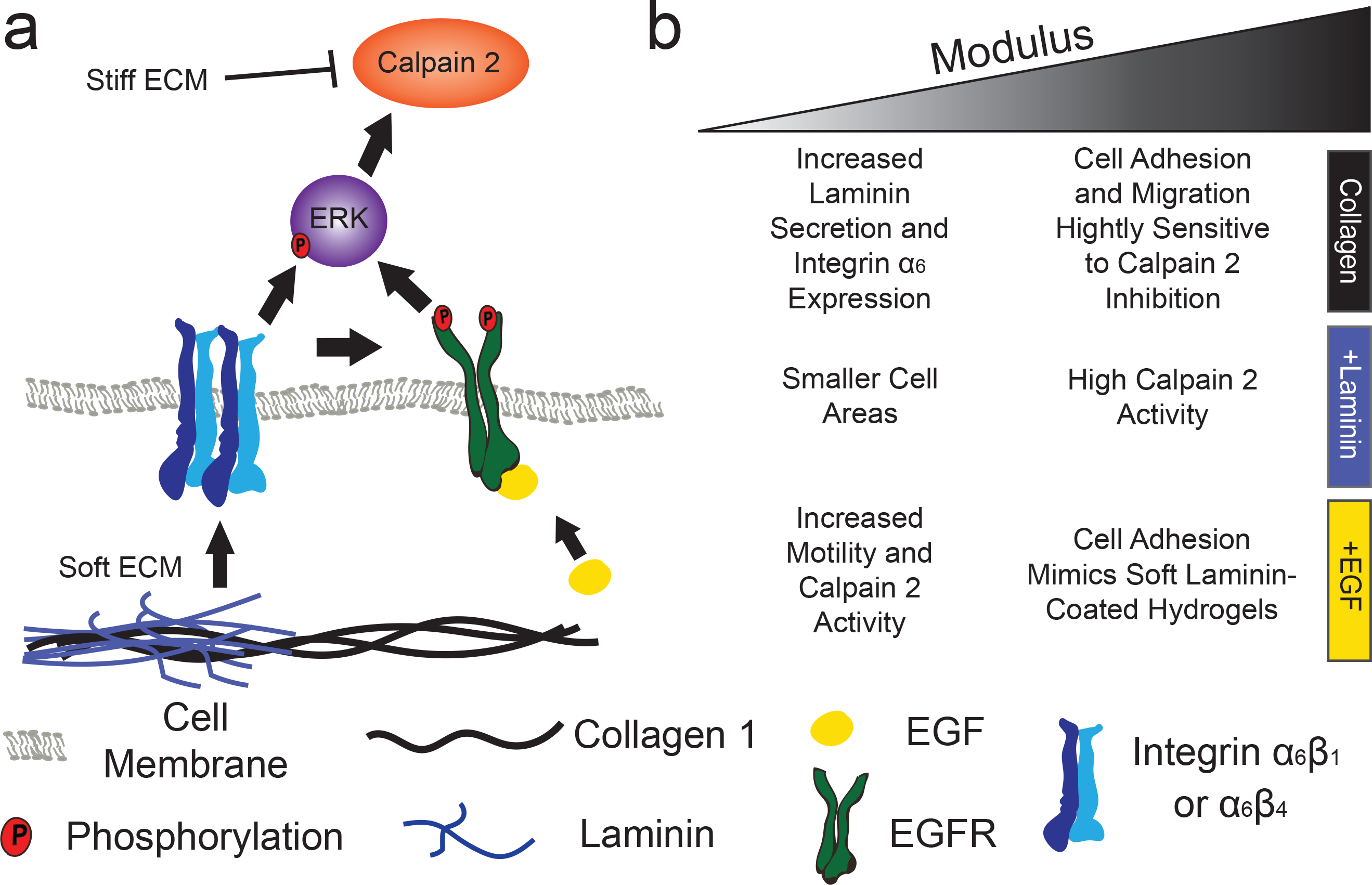
Mechanosensitive signaling is dependent on laminin and substrate stiffness. (a) Breast cancer cells on soft gels, or any stiffness gel supplemented with EGF, were small and motile. Cells on soft gels produced their own laminin to engage α_6_, or we added it exogenously. Our data suggests this increases activation of ERK and Calpain 2, which feeds back to increase turnover of focal adhesions. (b) During adhesion to substrates, cells were more responsive to laminin on soft substrates but did not adhere as well on stiff substrates with a calpain 2 inhibitor. Cell adhesion was facilitated in the presence of EGF on both soft and stiff gels. After 24 hours on gels, calpain 2 activity decreased with increasing stiffness, while EGFR phosphorylation increased with stiffness without added EGF. Cell motility had a biphasic dependence on substrate stiffness.

We focused this study on breast cancer cells because of the important roles for ECM binding and stiffness in metastasis. Cancer metastasis begins with individual or collective groups of cells migrating away from the primary tumor site [59-61]. This motility is driven in part by cell-ECM adhesion via integrins and the creation and turnover of focal adhesions, both of which are known to be sensitive to the stiffness of the surrounding ECM [62]. Integrins have well documented responses to changes in the microenvironment stiffness. As two examples, the expression of integrins α_2_, α_3_, and β_1_ is higher in mouse mammary epithelial cells cultured on TCPS compared to cells on a soft basement membrane-like matrix [50], and integrin α_5_ expression was found to increase 5-fold on a 10 kPa stiffness compared to 1 kPa in cancer cells [17] and over a similar range in fibroblasts [63]. Most of this work has focused on collagen- and fibronectin-binding integrins, with the exception of one paper on integrin α_6_ in fibroblasts, where expression increased with stiffness, resulting in increased invasion [64]. We demonstrate here the mechanosensitivity of integrin α_6_ in breast cancer cells, and the downstream regulation of cell adhesion, migration, and response to EGF (Fig. 1c, Fig. 2b). Integrin α_6_ is a laminin-binding subunit, and so many of the stiffness-sensitive phenotypes we observed either required, or were exaggerated by, addition of laminin to the culture substrate. However, cells exhibited mechanosensitive behavior on collagen alone when cells were on hydrogels for longer times. For example, we observed a biphasic relationship in migration speeds on collagen 1 only (Fig. 3e). We present data that on soft surfaces, cells produce extracellular laminin and engage integrin α_6_ in these instances, and that knocking down integrin α_6_ expression limited response to EGF (Fig. 2g).

Previous work has established the role of integrin α_6_ in key aspects of the metastatic cascade. Low integrin α_6_ expression inhibits the migration and proliferation necessary for establishing metastases at distant secondary sites [65]. Additionally, integrin α_6_ expression can protect cells from radiotherapy treatment both through PI3K and MAPK activity [66]. In a cohort of 80 patients, integrin β_4_ was co-expressed with integrin α_6_ and laminin in primary tumors, and co-expression of α_6_β_4_ with laminin production was significantly correlated with breast cancer relapse and death [43, 67]. This is further supported by other work in breast cancer, that elucidates the role of laminin-511 in cell adhesion and migration [18], which are key phenomena in the metastatic cascade. Our data demonstrate that this is a mechanoresponsive effect, as the most laminin is secreted by the breast cancer cells on the softest substrate (Fig. 1d). Interestingly, this phenomenon is not maintained in other secreted ECM proteins, such as collagen 1 (Fig. 1d) or fibronectin (Fig. S3a), suggesting a key role for integrin α_6_ binding to laminin on soft surfaces. This is possibly mediated through the mechanically responsive transcription factor, TAZ [68], which has been shown to regulate laminin production [69]. Here we found that both integrin β_4_ expression and laminin production increased after 6 days of culture on soft gels (Fig. 1b, Fig. 1d), where EGF enhances migration in an integrin α_6_-dependent manner (Fig. 2g).

We further observed that during cell adhesion, integrin α_6_ engagement with laminin decreased cell size and facilitated adhesion similarly to EGF (Fig. 2a-b). Smaller cells can be correlated with a cancer stem cell and invasive phenotype [70], and our work suggests integrin-mediated mechanotransduction contributes to that behavior. The α_6_β_4_ integrin dimer associates with EGFR [46], and we observed that integrin α_6_ engagement increased the sensitivity to EGF (Fig. 2b). Further, our data agrees with other work, which has demonstrated that the integrin dimer α_6_β_4_ plays an important role in cell response to growth factors, even in the absence of laminin [47]. We also observed that integrin α_6_ adhesion shows a similar phenotypic response on soft hydrogels as EGF stimulation.

Downstream of integrin α_6_ engagement and EGFR phosphorylation, cells activate calpain 2, which facilitates focal adhesion turnover, and therefore, motility (Fig. 3e-f). EGFR phosphorylation is necessary for mechanosensing [21] and activity in downstream effectors, including MEK, ERK, and calpain 2 at the cell membrane [71, 72], to cleave focal adhesion proteins vinculin, talin, paxillin, and FAK [73, 74]. Additionally, recent work has demonstrated that calpains are recruited to protrusions containing integrin β_4_ [75]. During adhesion, we saw that calpain 2 activity was higher in cells on the hydrogels compared to TCPS, which was consistent with RNA levels (Fig. 1a-b, Fig. S6c). Calpain 2 inhibition during adhesion resulted in larger, less motile cells on stiff substrates. The opposite trend in adhesion was observed when activating this pathway, as cells spread to a smaller area at both 1 and 41 kPa in the presence of soluble EGF (Fig. 2a). Together, these data suggest that EGFR phosphorylation could be a stronger activator of calpain 2 than stiffness. Previous work has shown that calpain 2 has a significant correlation with an epithelial-to-mesenchymal transition [76] and lymphatic or vascular invasion, where patients with basal-like tumors and high calpain 2 expression had a significantly worse prognosis [77]. Others have also found that this is an isoform-specific effect, as calpain 2, but not calpain 1, was upregulated in gastric cancer compared to healthy tissue [78]. Our work demonstrates a mechanosensitive role for calpain 2, downstream of both integrin α_6_ and EGF, providing further evidence for its potential as a druggable target for metastatic breast cancer.

## Conclusions

Mechanical forces have long been thought to play a role in tumor progression, and here we shed new light on two mechanoresponsive proteins that influence breast cancer cell motility: integrin α_6_ and calpain 2. First, enhanced integrin α_6_ expression on soft substrates is associated with increased adhesion and laminin secretion. Second, we find that both integrin α_6_ and EGFR activate calpain 2 in a mechanosensitive manner, which mediates cleavage of focal adhesion proteins necessary for motility. Others have suggested targeting calpains to inhibit cancer cell invasion [79], and here we provide a mechanosensitive role of calpain 2 using tunable hydrogel substrates.

## Author Contributions

ADS, SRP, and CCB participated in experimental design. ADS, CLH, and LEB performed experiments. All authors discussed results and contributed to editing of the manuscript.

